# The intercellular transfer of extracellular vesicles markers CD63, CD9 and CD81 is spatially polarized and restricted to cell vicinity

**DOI:** 10.64898/2026.02.23.707285

**Authors:** Marie Simon, Yé Fan, Hervé Acloque, Eric Rubinstein, Anne Burtey

## Abstract

Extracellular vesicles (EVs) are small lipid structures secreted by cells that originate from the cell surface (typically enriched in the tetraspanin (tspan) CD9) or from multivesicular bodies (typically enriched in the tspan CD63). Current methods for studying EVs involve concentrating and purifying EVs, without providing information about the distance or amount of EVs that may transfer from one cell to another. Here, we developed a coculture assay of human mammary MCF-7 cells to study the transfer of mCherry-CD81 or mCherry-CD9 from “donor” cells to a lawn of “acceptor” cells stained with cell tracker blue or green (CTB/CTG), non-transferrable fluorescent dyes. Using confocal fluorescence microscopy, we observed the presence of spots containing mCherry-CD81 or mCherry-CD9 outside donor cells, concentrated at short distance from donor cells and that overlapped with CTB signal, suggestive of their internalization in acceptor cells. Endogenous CD63, CD81 and CD9 also transferred more efficiently at short distances, even in the presence of a flow, as shown by immunostaining cocultures of wild type and KO CD-63, or −9, or −81 cells with antibodies directed against these tspans. Computation of the (x,y,z) coordinates of tspans-containing spots revealed a double polarized transfer: in (x,y), it distributed along a gradient that started from donor cells and decreased with the distance, and in (z), it was stronger in basal compared to upper planes, a (z) polarization that was affected by syntenin-1 depletion in donor cells. Simultaneous monitoring of CD9/CD81 transfer from into double CD81/CD9 KO cells showed that cells transferred more CD81 spots than of CD9. At the basal level, CD63 and CD81 spots were plasma membrane derived as they almost always contained CD9+, and resembled membranous remnants of migration. However, live cell imaging showed migration independent secretion of EVs in the extracellular space, in upper planes. Altogether, not only is our coculture assay suitable for the direct qualitative and quantitative study of EV-transfer, but it highlighted shared three-dimensional features of EV markers transfer between cells.

## Introduction

Extracellular vesicles (EVs) represent a novel principle of intercellular communication that has come into light in the last decades, with important role in physio-pathological processes. EVs are vesicles delimitated by a lipid bilayer secreted by most types of cells and that carry informative molecules to recipient cells. They represent a heterogeneous family of structures that were categorized in two main subclasses based on their composition, size and biogenesis mechanisms. On one hand, exosomes are structures of less than 150 nm in diameter that form in multivesicular endosomes (MVE) of the endocytic pathway. They are released in the extracellular space upon fusion of MVE with the plasma membrane and are enriched in CD63, a member of the tetraspanin (tspan) protein family. On the other hand, ectosomes are formed by evagination of the plasma membrane that pinch off in the extracellular space. They are enriched in molecules localized at the plasma membrane such as the tspan CD9 and its molecular partners^1,2^. Ectosomes are less limited in size, sometimes reaching up to several micrometers in diameter. Not only these EV subclasses differs in terms of their origin, mechanisms of formation, but also on their proposed function in several key processes such as cancer, immune response, neurological function^3,4^. The function of EVs may depend on their internalization in acceptor cells. For example, internalization of CD9-containing EVs was found to induce mitogen activated phosphor kinase (MAPK) cascade signaling that in turn favored cell migration and epithelial to mesenchymal transitioning of pancreatic cells^5^. However, the sole deposition of EVs on cell culture vessels favored mammary breast cancer cells MCF-7 migration^6^. EV release in the extracellular space, displacement, docking on cell surface and/or internalization, is a process that we referred to here as “EV transfer”. In the rare studies that addressed the transfer of EVs, authors have used experimental approaches that involved substantial processing of EVs (concentration, purification, fluorescence labelling) prior addition onto acceptor cells^7^. Not only it is difficult to exclude that EV processing did not interfere with their structure and docking abilities, but it is also difficult to interpret the relevance of their detection in acceptor cells given the considerable number of EVs added. It also remains unclear how many EVs are transferred from one cell to another, in physiological settings, or at least without extensive EV processing. In the zebrafish embryo, exogenously expressed fluorescently-tagged CD63 was detected in the general circulation and in non-expressing cells located at distance from expressing cells^8^, suggesting that *in vivo*, fluorescent-tagged CD63 may have travelled long-distance provided their secretion in the general circulation. Here, we developed an assay to directly monitor EV transfer without EV processing, a platform to screen for molecular regulators of EV transfer. Based on this approach, we studied the 3-D transfer of the three main EV markers: CD63, CD9, CD81, either exogenously expressed as mCherry-fusions or endogenously expressed. We provide clues on the (i) the rate of transfer, (ii) the number of EV markers containing spots that donor cells transferred, (iii) the distance at which they travelled from donor cells, (iv) the influence of the flow and of syntenin-1 in regulating the topology of EV transfer in the three dimensions.

## Results and discussion

### A coculture assay to analyze the transfer of EVs: m-Cherry tagged tspans

To study EV transfer, we adapted a coculture assay previously developed to analyze tunneling nanotubes^9^. As a cellular model, we used the mammary cell line MCF-7 for which the production of exosomes has been previously characterized^10^. MCF-7 “acceptor” cell population was labelled with non-transferrable dyes (cell tracker blue (CTB) or green (CTG)) prior coculture with a “donor” MCF-7 cell population that we first transfected with EV markers tagged with mCherry (Fig. 1). We first analyzed the transfer of CD9, marker of plasma membrane (PM)-derived EVs, and of CD81, another tspan localized at the plasma membrane in MCF7 cells^2^, used as a marker of small EVs^1,3^, and detected in CD9 positive EVs from different cell types^3,4^ or in human plasma^3^. Cocultures were done at low donor:acceptor (1:400) cell ratio and high confluence to restrict cellular movements, with cell-cycle blocked to avoid overcrowding and dilution of the plasmid in daughter cells as we have previously done^9,11^. By doing so, the transfer of EVs could be monitored directly by 3D high resolution confocal microscopy, from donor cells that were surrounded by many acceptor cells. Confocal stacks of images were centered on donor cells respecting criteria defined in ^9,11^: exclusion of small, rounded cells (attachment failure or cell death), multinucleated cells, and subconfluency areas. After 48 h of coculture, we observed that nearly all donor cells transfected with either mCherry-CD81 or mCherry-CD9 were surrounded with fluorescent “spots”, demonstrating the release of material positive in these markers, presumably EVs (Fig. 1A, black and pink arrowheads and S1A for the quantification of the percentage of donor surrounded by fluorescent spots). As a control, very few and dim fluorescent spots were detected around fewer cells expressing mCherry only (Fig. 1A “mCherry alone”, and S1A).

**Figure 1:**
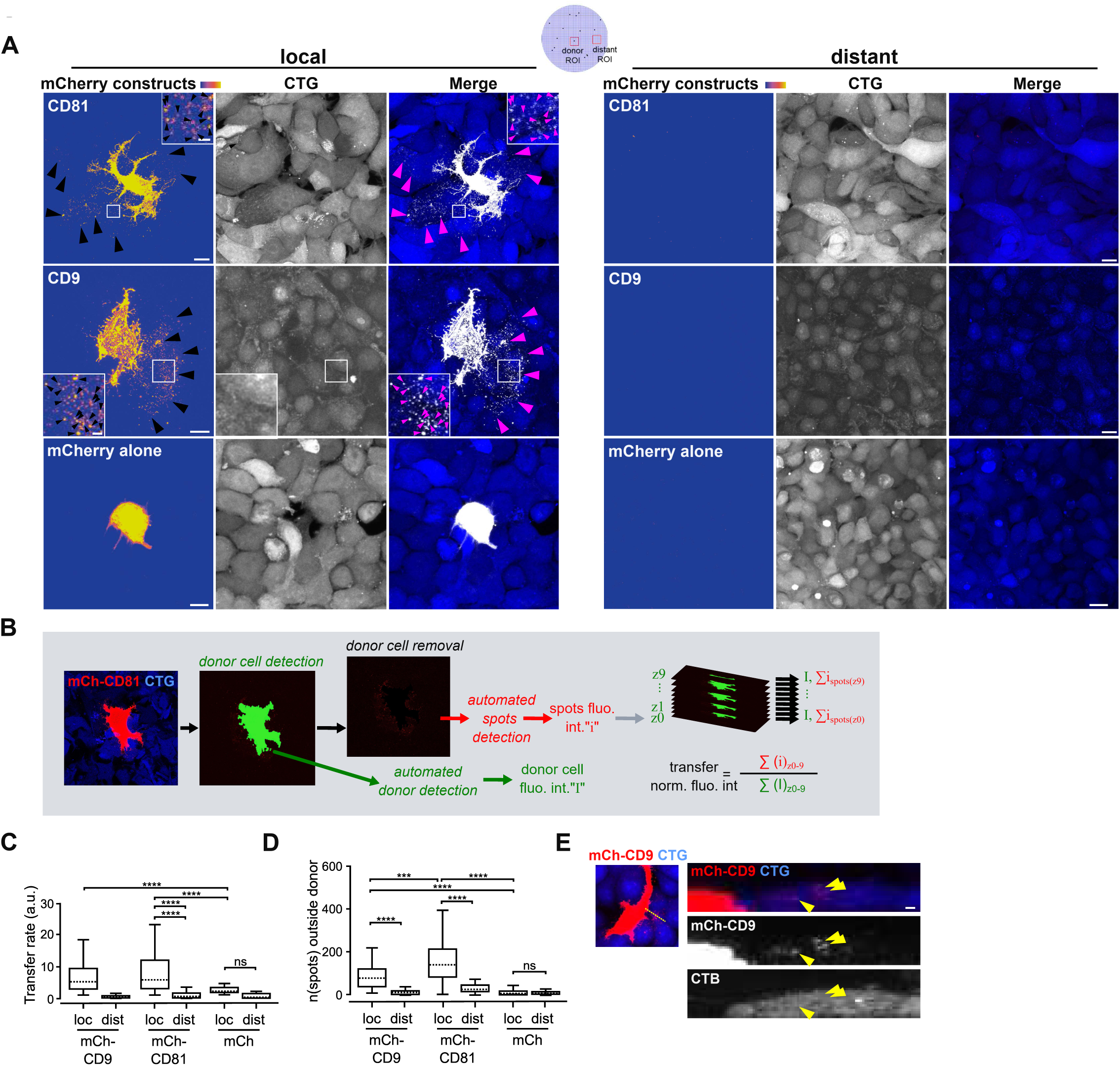
Excretion of mCherry-CD9, mCherry-CD81, or mCherry alone from donor MCF-7 cells to adjacent MCF-7 cells. MCF-7 cells transiently expressing mCherry-CD9, mCherry-CD81 or mCherry alone were cocultured for 48 h with non-transfected MCF-7 cells stained with Cell Tracker Green (CTG). After fixation and mounting, cocultures were imaged by confocal microscopy. **A.** Representative maximal z-projections of stack of confocal images acquired in regions of the cocultures containing acceptor cells stained with CTG (in blue) surrounding a donor cell (“local”) or distant from donor cells (“distant”). Ametrine LUT colormap is used to visualize mCherry. Arrowheads point to mCherry-positive spots outside donor cells. Scale bars: 10µm. Insets are magnifications of ROIs, scale bars: 2µm. **B.** Overview of the ICY software workflow. **C**. The rate of transfer is given as the summed mCherry fluorescence in spots detected outside the donor cell normalized to the donor cell total fluorescence. **D.** The number of spots detected outside donor cells. In C and D, data are shown as boxplots (median, interquartile range, whiskers, and outliers) from at least three independent experiments. Number of stacks analyzed: mCh-CD9: n= 53, mCh-CD81: n= 48, mCherry alone: n= 56 for local regions; mCh-CD9: n= 17, mCh-CD81: n= 18, mCherry alone n= 19 in distant regions. **E.** Reslices along a line of interest (dotted yellow). Arrowheads point to transferred mCherry-CD9 spots. Scale bar: 1µm. **F.** Percentage of spots containing CTG.

As shown in Figure 1, the fluorescent spots observed outside donor cells appeared to be more abundant at a closer distance to donor cells than in control regions that were randomly selected at distances of at least 8 cell diameters away from any donor cell (Fig 1A, compare “local” *vs*. “distant”). To quantify the transfer, we developed a protocol on ICY software^12^. All stacks of images were acquired with the same settings consisting in ten confocal planes covering the total donor cell height, from top to bottom, with a pixel size of 140 nm in (x,y). For each stack, the workflow (i) detected the donor cell, (ii) measured its volume and fluorescence level, and (iii) removed the donor cell after enlargement of its contour by 4 pixels to avoid the artefactual detection of cellular protrusions as transfer, (iv) detected spots of mCherry fluorescence (v) and measured the fluorescence intensities inside spots and the number of the latter (Fig. 1B). The transfer rate was defined for each stack as the sum of the fluorescence intensities in spots normalized to that of the donor cell (Fig. 1B). Results of this analysis confirmed that there was more fluorescence intensity (Fig. 1C) and spots (Fig. 1D) at local positions than at distant positions (Fig. 1C-D “loc” *vs* “dist”). It further revealed differences between CD9 and CD81. On one hand, the intensity measured outside donor cells was 14 times higher in local *vs.* distant regions for cells transfected with mCherry-CD9, compared to 8 times higher for cells transfected with mCherry-CD81 (Fig. 1C). This difference is likely due to a stronger signal of mCherry-CD81 at distant locations (Fig. 1C, left graph, “distant”, in which the values of the 3sd quartile were equal to 1.48 for mCherry-CD81 and 0.82 for mCherry-CD9). On the other hand, when considering the number of spots detected, donor cells were surrounded by an average of 181.0 (+/- 21.8) mCherry-CD81 positive spots compared to 101.0 (+/- 13.2) mCherry-CD9 spots and only 20.0 (+/- 4.9) of mCherry alone spots (Fig. 1D). Importantly, these numbers are in the same order of magnitude than the number of EVs detected in the medium of the same cells (200 EVs/cell)^2^. The higher detection of CD81 spots compared to CD9 spots is not due to a higher expression level of mCherry-CD81 compared to mCherry-CD9, as shown in Fig. S1B. This is further confirmed by the finding that the transfer rate was similar for both mCherry-CD9 and mCherry-CD81-transfected cells, and that the number of spots was independent of the cell expression level in the donor cell (Fig. S1C-D). This may therefore be explained by a higher production/release of CD81-expressing EVs, which is in line with previous findings showing that in several cell lines, including MCF7, CD81 is more enriched in EVs than CD9^2,13^. Alternatively, CD9-positive EVs could be less able to bind to cells than CD81-positive EVs, or be faster internalized and degraded leading to the subsequent dilution of the CD9 signal and thereby lesser CD9 spots detected. To get insights in the internalization of EV markers, we took advantage of the fact that CTB and CTG are cytosolic dyes^9,11^ and reasoned that if EVs were internalized in acceptor cells, the fluorescence signal in spots should be overlapping with that of CTB/CTG and on the other way around, non-internalized EV markers should be localized in CTB/CTG low signal regions. Results showed overlapping of EV marker positive spots with CTB signal on single confocal planes suggesting that some EV markers were perhaps internalized (Fig. 1E, arrowheads).

### A coculture assay to analyze the transfer of EVs: endogenous tspans

To exclude the possibility that addition of the tag or over-expression affected the transfer of the tspans, we analyzed the transfer of endogenous EV markers: CD81, CD9 but also CD63, the consensus markers of exosomes^1,3^. To that end, we cocultured wild type MCF-7 cells with knocked-out (KO) CD9, CD81 or CD63 MCF-7 cells described previously^2^. The transfer of endogenous proteins was studied by immunostaining with antibodies directed against CD9, CD81 or CD63 revealed by secondary fluorescent antibodies (Fig. 2A-B). Consistent with the results observed for fluorescent proteins, the number of spots was for all markers in the same order of magnitude as the number of EVs collected from these cells (Fig. 2C) and the transfer was more efficient locally than at distant positions (Fig. 2B-C). A difference with the above experiment, is that the number of CD81-positive spots and of CD63 spots, but not that of CD9-positive spots, correlated with the intensity of cell labeling for the majority of cells having the highest level of expression (>15.10^8^ a.u., Fig. S2A).

**Figure 2:**
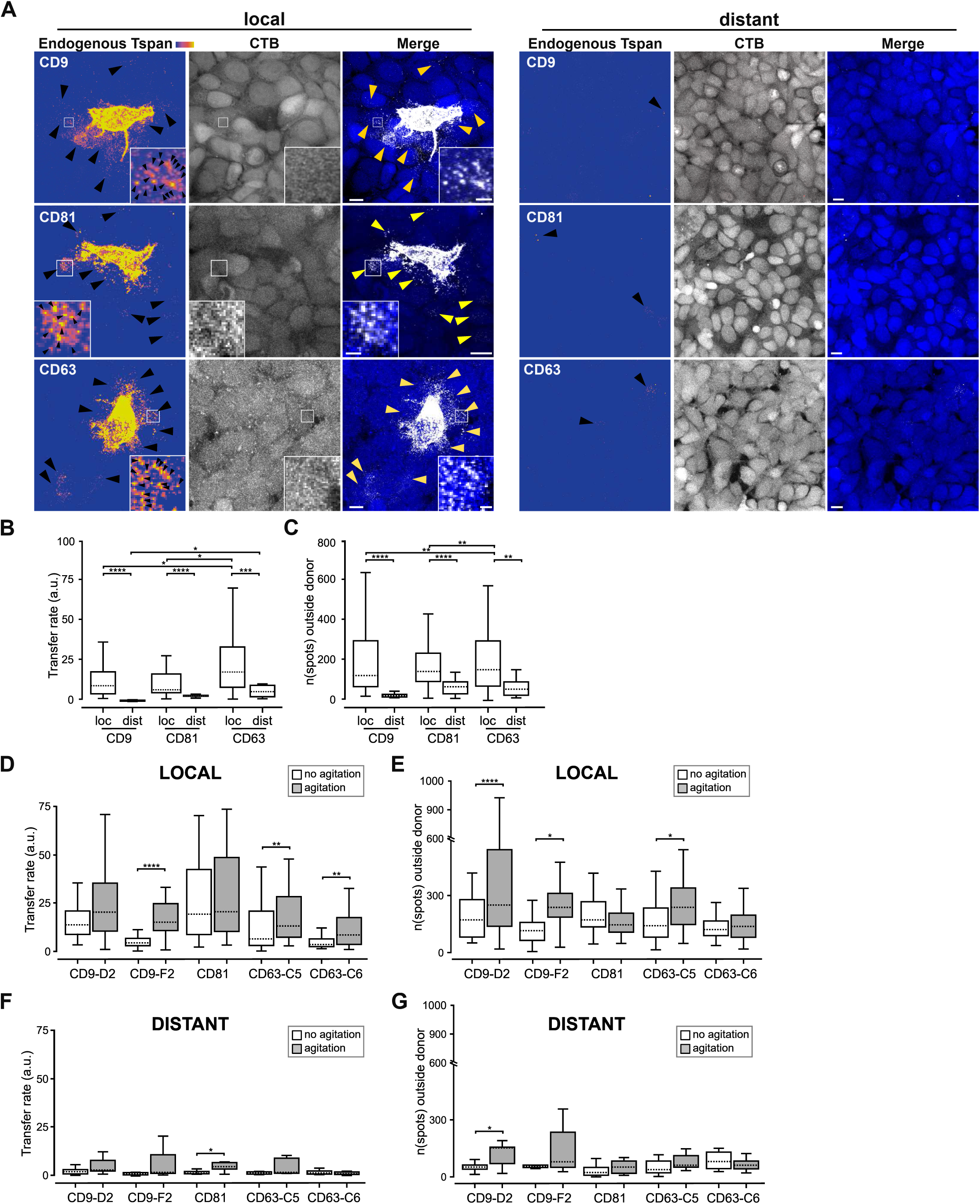
Excretion of endogenous CD9, CD81 or CD63 from single donor MCF-7 cells to adjacent KO cells. MCF-7 wt cells were cocultured for 48 h with MCF-7 cells KO for CD9, CD81, or CD63 stained with Cell Tracker Blue (CTB). Cells were fixed, permeabilized and immunostained against CD9, CD81, or CD63 revealed by a secondary antibody F(ab)^2^-AF555 prior imaging under a Leica SP5 confocal microscope. **A.** Representative maximal z-projections of confocal planes of regions of the cocultures surrounding a donor cell (“local”) or distant from donor cells (“distant”). CD9, CD63 and CD81 channels are displayed using ametrine LUT colormap for better visualization, or in white in merged images. Arrowheads point to positive spots outside donor cells. Scale bars: 10µm. Insets show magnifications of ROIs, scale bar: 2 µm. **B.** Rate of transfer **C.** Number of spots outside donor cells. **D-G.** Transfer rate or number of spots in the presence (grey boxes) or absence (white boxes) at local or distant sites. Number of stacks analyzed: CD9: n= 52, CD81: n = 50, CD63: n = 42 in local regions, CD9: n = 9, CD81: n = 14, CD63: n = 12 in distant regions.) D-G. **D-G.** Transfer rate (D-F) or number of spots (E-G) in the presence or absence of agitation at local (D-E) or distant (F-G) sites. Number of stacks analyzed: CD9-D2: n= 41, CD9-F2: n = 39, CD81: n = 41, CD63-C5: n = 39, CD63-C6: n = 36 in local regions under agitation, CD9-D2: n = 42, CD9-F2: n = 40, CD81: n = 41, CD63-C5: n = 43, CD63-C6: n = 29 in local regions under no agitation, CD9-D2: n = 9, CD9-F2: n = 8, CD81: n = 9, CD63-C5: n = 9, CD63-C6: n = 8 in distant regions under agitation, CD9-D2: n = 9, CD9-F2: n = 9, CD81: n = 10, CD63-C5: n = 9, CD63-C6: n = 8 in distant regions under no agitation.

Comparing the transfer rate and spot number per experiment suggested that there was some variability according to the experiments (data not shown). This could depend on the variability of donor cells but also on acceptor cells. To further study this variability, we analyzed the transfer of EV markers towards two clones of CD9 KO and CD63 KO cells. Figures 2D and 2E, showed the existence of an acceptor clone-dependent efficiency of the transfer since the rate of transfer, and the number of spots transferred, were different for the two CD9 or CD63 clones (Fig. 2D-E, white boxes). For each clone, the variability remained high suggesting some variability due to the acceptor cell. As stated for CD81 and CD63, the expression level of the donor cells contributed to this variability.

### Agitation poorly stimulates long range transfer

The above observation showed that only a few spots were detected at distance from donor cells whether we studied endogenous tspans (Fig. 2F-G) or mCherry-tspans (Fig. 1C). We hypothesized that this might be due to the lack of flow in the culture. To address this question, cells were cocultured under agitation. Results showed that despite agitation, the CD9-, CD63 and CD81-positive spots released by donor cells remained mainly local (Fig. 2D-E, grey boxes) when compared to distant sites (Fig. 2F-G, grey boxes). In these distant sites, agitation neither modified the transfer rate (Fig. 2F) nor the number of spots (Fig. 2G). However, agitation did increase the transfer rate locally, surrounding donor cells, in the low acceptor CD9 and CD63 clones (Fig. 2D, compare white *vs*. grey boxes). For CD63-C6 clone, agitation increased the transfer rate but not the number of spots (Fig 2D vs. 2E), as if agitation stimulated the number of CD63 molecules in EVs. These results suggested that the flow was not sufficient to drive long-distance transfer in this system. This could be due to the dispersion of EVs by agitation and/or agitation that prevented the docking of EVs on acceptor cells, and therefore detection. When it came to the increased local transfer upon agitation, it is in line with the increased yield of EV produced when applying a turbulent flow on cell cultures^14,15^.

### EV Transfer mainly occurs to adjacent cells

The above data showed minimal transfer of the three tspans studied at long distance. Closer observation of our images, as those shown in Fig 1, suggested that transfer occurred mainly to the cells that were adjacent to donor cells, and not to those located a few microns away, suggesting that EV may transfer to even shorter distances than 8 cell diameter (*i.e.* the minimal distance of distant regions) (Fig. 3A). To validate and quantify this, we computed the (x,y) coordinates of each spot detected outside donor, and their position relative to the closest border of the donor cell (Fig. 3A-B). The distribution of spots in each plane of each field acquired (160 µm x160 µm field size) was classified according to their distance from the donor into four distances ranges (0-20 µm, 20-40 µm, 40-60 µm and >60 µm) (Fig. 3C). It revealed that most of the spots outside donor cells were located at a distance inferior to 20 µm away from donor cell, and showed the existence of a gradient, that varied slightly depending on the tspan considered (Fig. 3C), a gradient that was not observed in distant regions (Fig 3C, “distant”). Whereas CD9 gradient was marked, it was more gradual for CD63-C6, the latter appearing to spread further away (Fig. 3C, green vs. pink). The distribution of spots was then analyzed in acceptor cell clones and upon agitation. The results confirmed that most of the transfer occurred at very short distances and that agitation has a low to null impact on the transfer. Interestingly, the transfer of CD63 to the low acceptor clone (C6) was stimulated by agitation and was associated with a stronger transfer at short distance.

**Figure 3:**
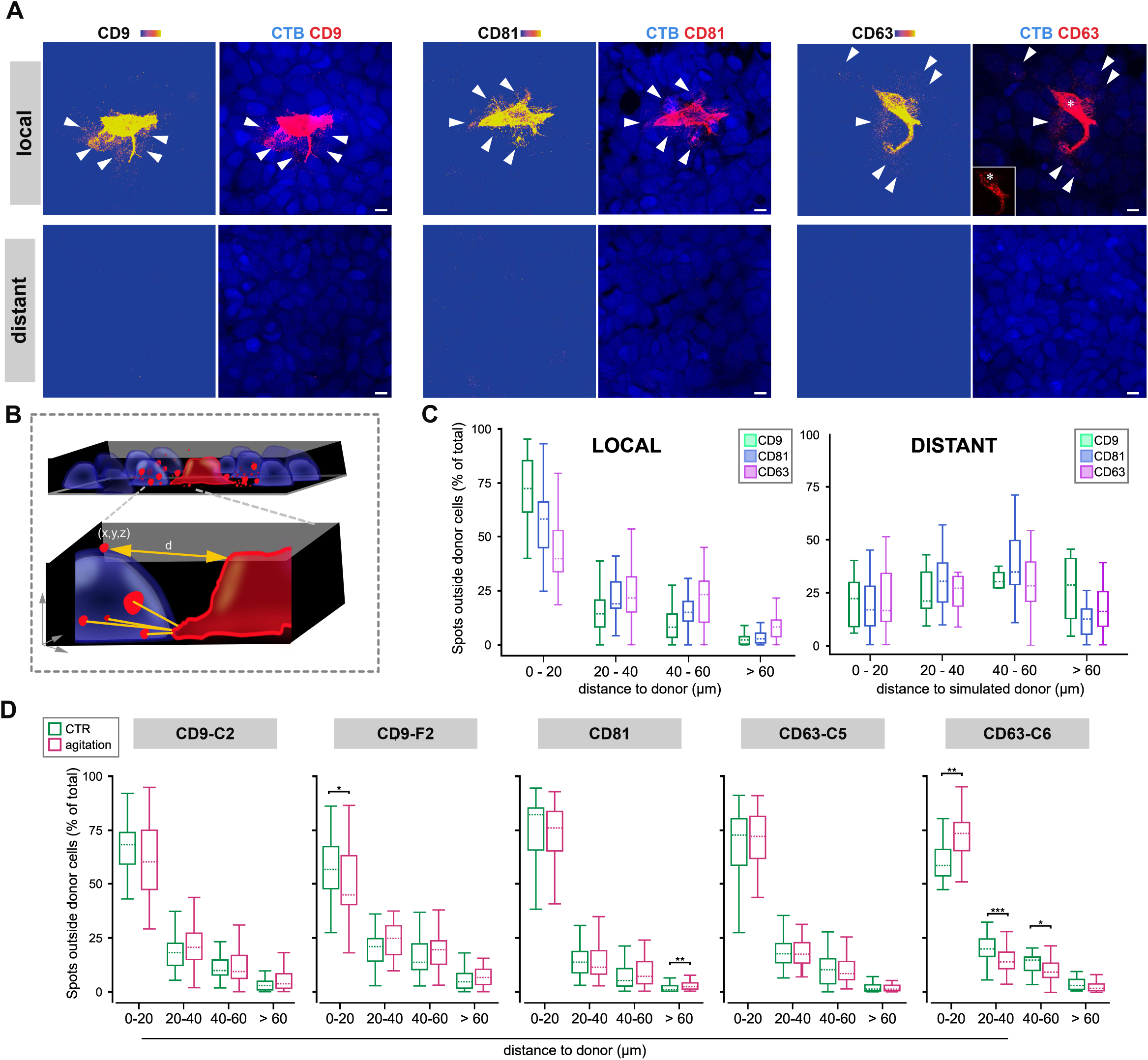
Distance of transfer along the (x,y) axis for endogenous CD9, CD81, or CD63. **A.** Representative maximal z-projections of confocal planes of regions of the cocultures containing acceptor cells stained with CTB (in blue) surrounding a donor cell. Arrowheads point to transferred spots. Scale bars: 10µm. **B.** Schematic representation of the (x,y,z) coordinates determined for each spot and the shortest distance from donor cell plasma membrane **C.** Percentage of spots located outside donor, at a given distance from the donor cell (0-20 µm, 20-40 µm, 40-60 µm, >60µm) and at distant regions from an artificial cell. **D.** Percentage of spots at given distances in the presence (red) or absence (green) of agitation.

### Distribution of EVs according to the vertical axis

Next, we analyzed the position of transferred spots along the (z) axis. Surprisingly, we observed a “polarization” of the transfer characterized by the accumulation of many spots at the substrate level, with or without acceptor cells “sitting” on top (Fig. 4A, “single basal plane”). These basal spots were observed for CD9 (Fig. 4A), CD63 (Fig. 4A) and CD81 (not shown). The number of spots in each of the ten planes stacks were automatically counted and plotted (Fig. 4B-C). Results showed that there were more spots in the basal planes than in upper planes surrounding donor cells (Fig. 4B), and for CD9, these basal spots represented nearly half of the spots (Fig. 4B). Regarding CD63, the number of spots in basal levels may be underestimated, as basal spots were often detected as part of CD63 donor cells and not taken into account by our automated protocol. Of note, that very few spots were detected at apical position in Fig. 4B, could be due to the washing steps of the immunostaining protocol. At distant sites, where the number of basal spots was low, no tetraspanin exhibited a higher proportion of basal spots. (Fig. 4C). Polarization of the transfer was further confirmed by plotting the distribution of spots in basal *vs*. summed upper *vs* summed apical planes (Fig. 4D). To understand whether this baso-apical distribution was also polarized along the (x,y) axis, *i.e.* distributed in a gradient, we plotted the vertical positions of the spots (basal *vs*. middle) according to the distance from the donor cell. Results showed that basal spots were mostly detected at distances inferior to 20 µm from donor cells, but in middle planes, more spreading was observed, particularly for CD63 (Fig. 4E).

**Figure 4.**
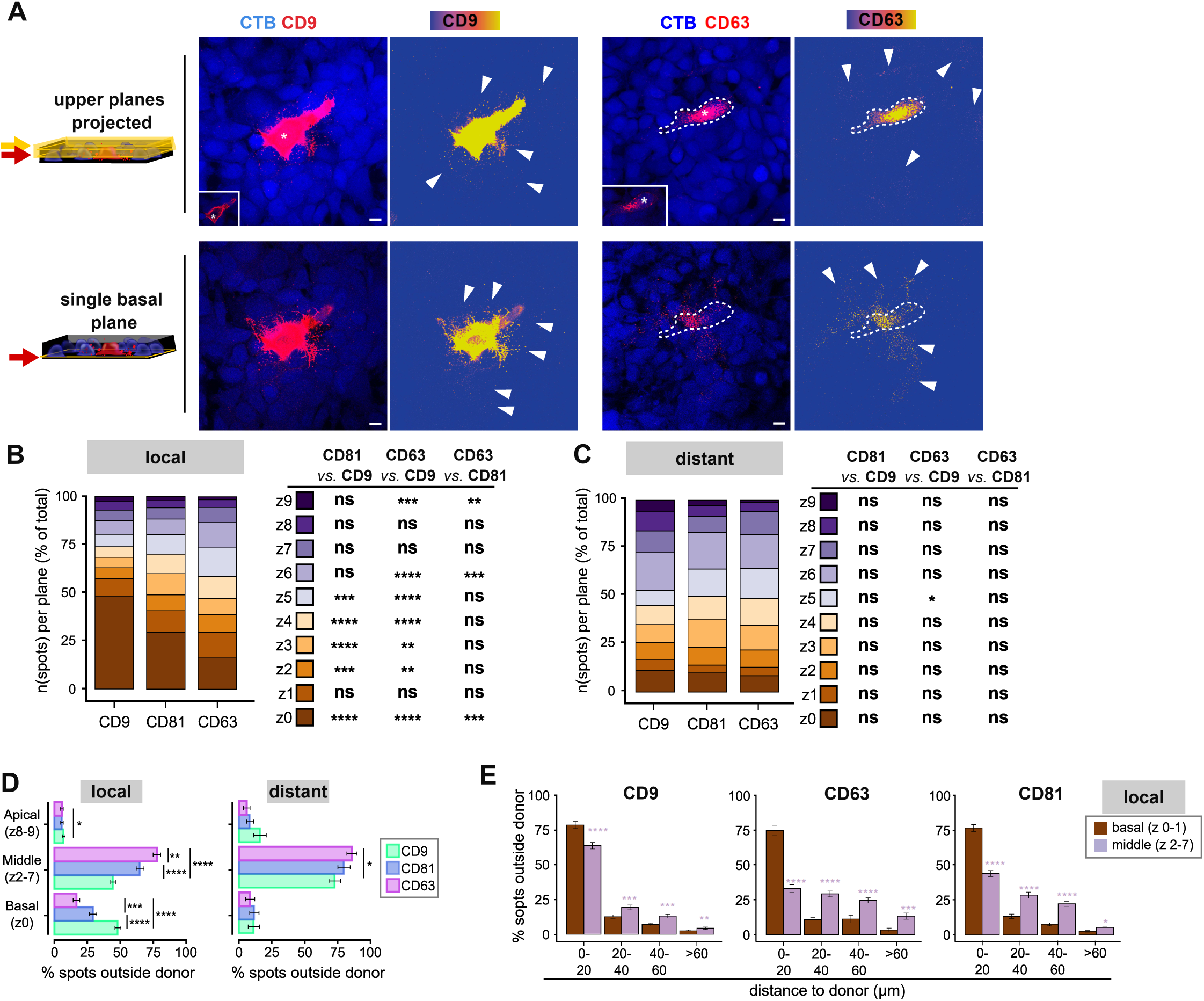
Localization of transferred CD9, CD63 and CD81 along the (z) axis. **A.** Representative images of single basal plane (lower panel), and of a maximal projection of upper planes (2-6.8 µm) of MCF-7 cells cocultured with KO CD9 cells immunostained with anti-CD9 antibody, and MCF-7 cells cocultured with KO CD63 cells immunostained with anti-CD63 antibody. Stacks consisted in 20 planes (for CD9) and 25 planes (for CD63). Scale bars: 10µm. Insets show donor cells at low intensity illumination **B-C.** Distribution of positive spots outside donor cells located on individual confocal planes as a percentage of total spots in all planes summed at local (B) or distance (C)**. D.** Percentage of positive spots outside donor cells located on basal, summed middle, and summed apical planes. **E.** Percentage of positive spots on basal or summed middle planes, according to the distance from donor.

### Dual monitoring of EV marker transfer

So far, our results strongly demonstrated that the transfer of CD9, CD81 or CD63-positive EVs is more efficient at short distances and showed some variability in terms of overall transfer, number of spots or distribution along the distance, preventing appropriate comparison between different tspans. This led us to compare the transfer of CD81 and CD9 by coculturing wild-type MCF-7 with double KO (dKO) cells for CD9 and CD81 expression (Fig. 5A). Immunostaining against CD9 and CD81 revealed interesting differences. CD81 transfer was higher than CD9 in terms of intensity and spot number (Fig. 5B-C). This was observed using the two clones of dKO acceptor cells, one having a more pronounced phenotype than the other (Fig. 5B-C). CD81 and CD9 transfer occurred mainly to the first 20 µm from donor cells (Fig. 5D), and analysis along (z) axis showed that a higher proportion of CD9 spots was located at basal position compared to CD81 (Fig. 5E-G, and S2D). Analysis of their relative localization outside donor cells was done on confocal stacks acquired at the highest resolution in (x,y) (Fig. 6A) showed their colocalization in single confocal planes (Fig. 6A-B). Interestingly, more of them (70 %) colocalized in basal spots (Fig. 6A “z=0”, 6C) than in upper spots (50%, Fig. 6D). This suggested that CD9/CD81 double-positive spots (presumably EVs) behave differently than single positive spots. The labeling of acceptor cells by CTB allowed an analysis of the internalization of the tetraspanins-positive structures that showed nearly all basal spots colocalized with CTB but considering the resolution of light microscopy, we do not conclude whether they were internalized or not in basal level (Fig. 6C, reslice along line 2). In contrast, in acceptor cell upper planes, we detected spots containing either CD9 or CD81, or both, not superimposing to CTB (Fig. 6A, z= 3.25 µm, inset, arrowhead and 6C, reslices along lines 1-2), which could correspond to non-internalized EVs or EVs internalized but still in early endosomes containing extracellular medium, hence CTB^-negative^. Consistent with the automatic detection of Fig 6D, CD81 and CD9 did not always colocalize in upper planes (Fig. 6B), as also showed by reslicing of stacks in which apical spots were also detected (Fig. 6C).

**Figure 5:**
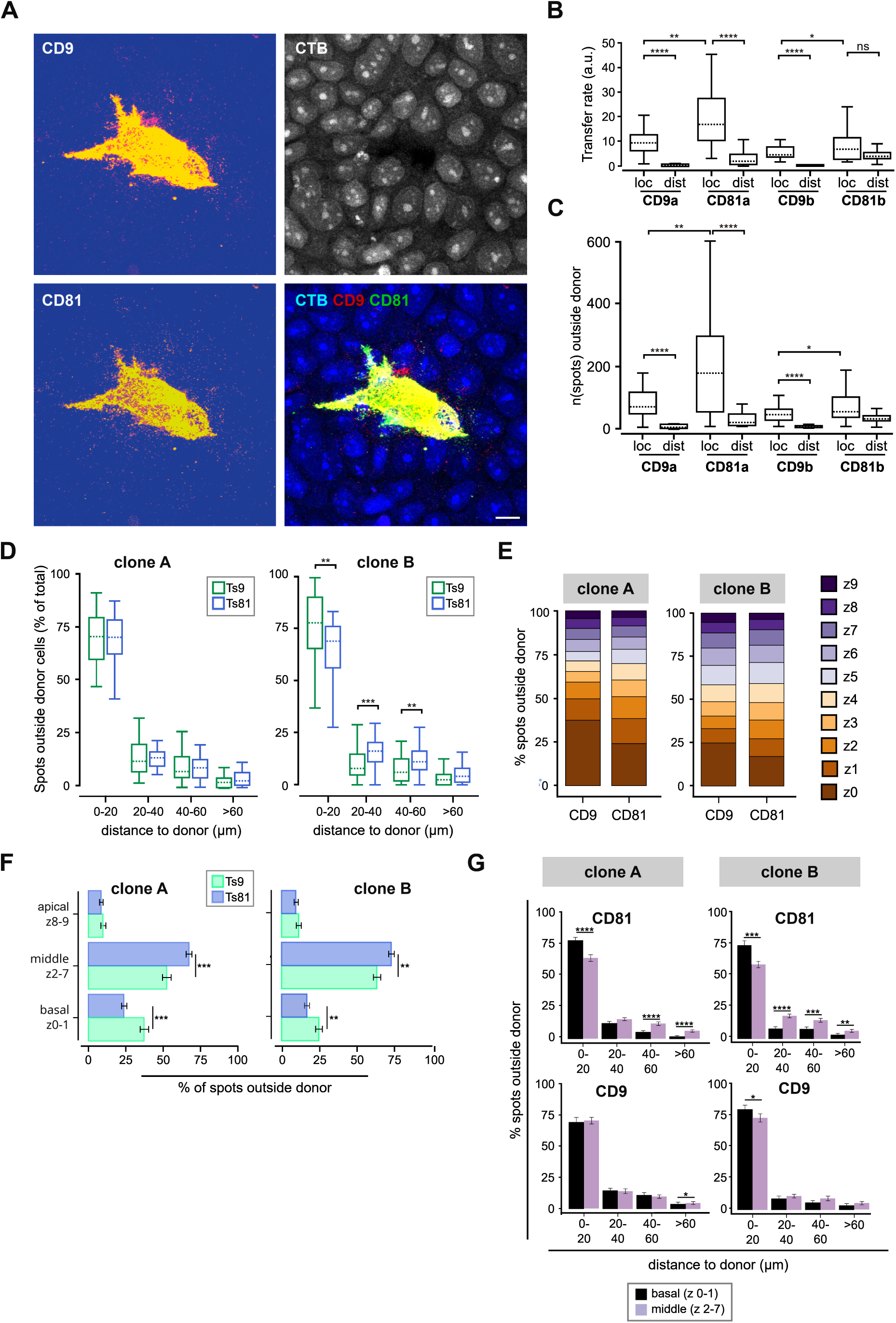
Dual monitoring of endogenous CD9 and CD81 transfer to double KO MCF-7 cells. MCF-7 cells were cocultured for 48 h with MCF-7 cells KO for both CD9 and CD81 (clone A or clone B) stained with CTB, fixed and immunostained with mouse anti-human antibodies CD9 (Mm2/57) and CD81 (5A6) revealed by subclass specific secondary antibody anti IgG1 coupled to AF568 and anti IgG2a coupled to AF488. **A.** Representative single confocal planes of regions of the cocultures with donor cells (ametrine/white) and acceptor cells stained with CTB (in blue). Scale bar: 10µm. **B.** Transfer rate. **C.** Number of spots detected outside the donor cell **D.** Percentage of spots according to the distance from donor cells. **E.** Percentage of positive spots outside donor cells on individual confocal planes (z0–z9)**. F.** Percentage of positive spots outside donor cells located on basal, summed middle, and summed apical planes. **G.** Percentage of positive spots on basal or summed middle planes, or summed apical planes according to the distance from donor, with or without agitation. Number of stacks analyzed: clone A: n= 35, clone B: n = 41 in local regions, clone A: n = 10, clone B: n = 13 in distant regions)

**Figure 6.**
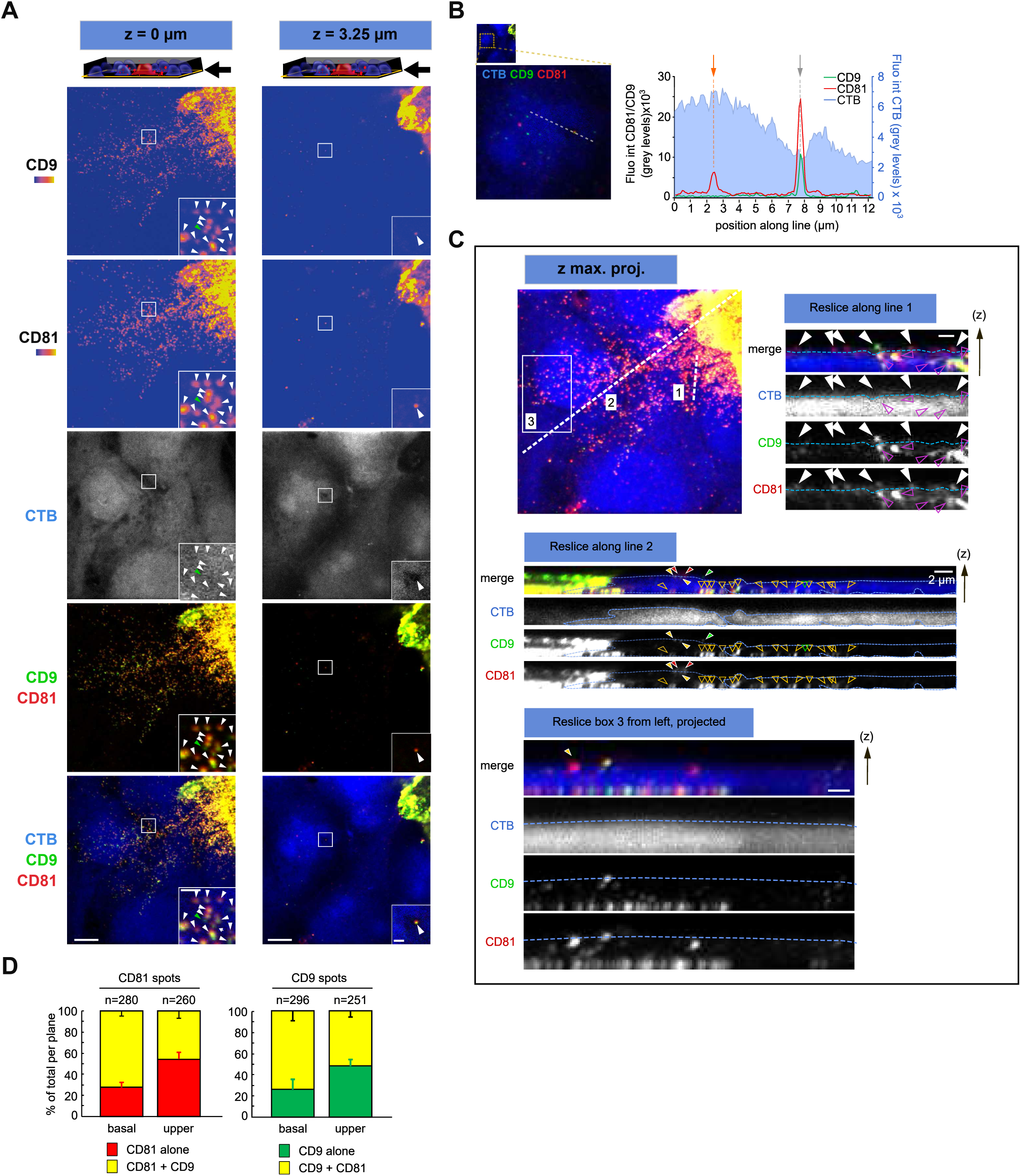
Colocalization of CD9 and CD81 upon secretion relative to CTB. MCF-7 cells were cocultured for 48 h with MCF-7 cells KO for both CD9 and CD81 (clone A or clone B) stained with CTB, fixed and immunostained with mouse anti-human antibodies CD9 (Mm2/57) and CD81 (5A6) revealed by subclass specific secondary antibody anti IgG1 coupled to AF568 and anti IgG2 coupled to AF488. **A.** Representative single confocal planes of regions of the cocultures at the interface donor/acceptor on high resolution images (80nm pixel size). Scale bar: 5 µm (1µm in insets). **B.** Intensity profiles of CD9, CD81 and CTB along a line of interest that crosses two spots, one CD81^+^CD9^-^ and the other CD81^+^CD9^+^, both CTB^+^. **C.** Maximal projection of the stack showed in A. Reslices along ROIs: line 1 (upper right), line 2 (middle) and box 3 (sliced from left, then maximally projected, lower panel) are shown. Scale bars: 1µm unless indicated otherwise. **D.** Quantification of the fluorescence intensity of CD9 in CD81 spots and vice versa, in basal or upper planes, normalized to background.

To compare the distribution of CD9-positive and CD63-positive spots, and to confirm previous observations in living cells, we analyzed living MCF-7 cells transfected with a bicistronic plasmid expressing GFP-CD63 and mCherry-CD9 and cocultured with CTB wild-type cells (Fig. 7A). In living cells, the transfer was local, oriented, and polarized in (z), for both tspans (Fig. 7A, arrowheads), the latter confirmed by quantification of the number of spots in the different planes (Fig 7B). To quantify the colocalization, mCherry-CD9 spots were automatically detected and the fluorescence intensities of GFP-CD63 was measured. The percentage of CD9 spots containing CD63 is shown in Fig. 7C (pink bars) and indicated that 40% of basal CD9 spots also contained CD63 but on the other way around, automated detection of GFP-CD63 spots showed that all basal GFP-CD63 spots contained mCherry-CD9 (Fig. 7C, green bars). These results suggested that basal spots are likely to derived from plasma membrane, given the omnipresence of CD9 in these, and that CD63 basal spots are also likely to be plasma membraned-derived EVs, given their content in CD9.

**Figure 7:**
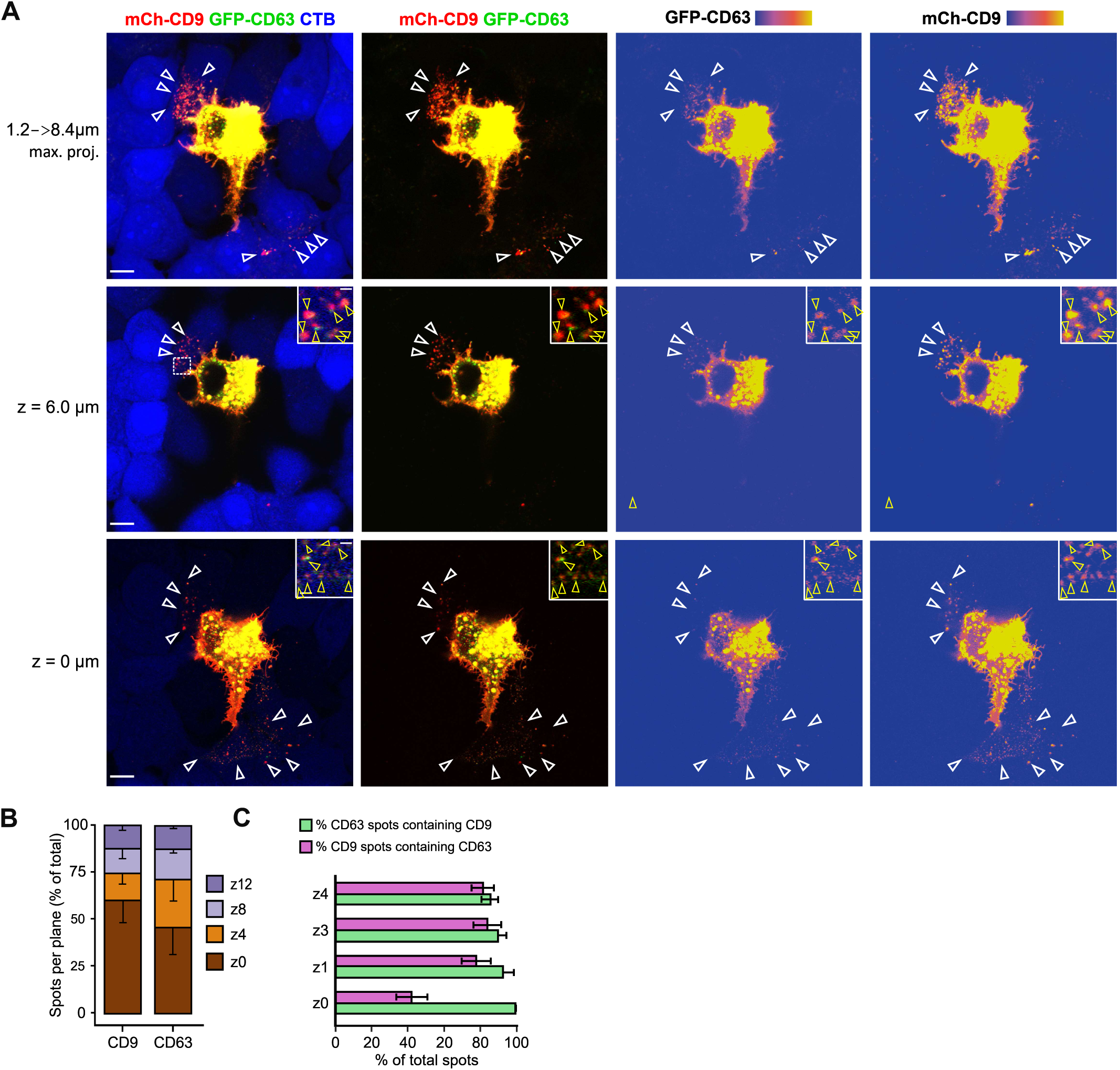
Live cell imaging of mCherry-CD9 and GFP-CD63 transfer simultaneously. MCF-7 cells transiently expressing bicistronic plasmids encoding for mCherry-CD9 and GFP-CD63 were cocultured with non-transfected MCF-7 cells, left for 4 h to adhere and imaged overnight every 15 minutes. **A.** High resolution confocal image showing the transfer of mCherry-CD9 and GFP-CD63 in basal (z=0), middle (z=6µm) or upper planes maximally projected. Open arrowheads point to transferred spots. Insets are magnifications of regions of interest. Scale bars: 10µm (in insets 1µm). **B.** Average number of spots in 4 confocal planes, as a % of total spot number of the plane. **C.** Average number of spots single or double positive in 4 confocal planes, expressed as a % of total spot number.

### Polarized released and cell uptake of EVs

We observed that the distribution of CD9, CD81 or CD63 spots in basal planes was often oriented towards one or a few adjacent cells (Fig. 8A), instead of being evenly distributed all around donor cells (Fig. 8B). Quantification of the number of donor cells presenting a transfer to only one side of donor cells showed that this oriented transfer was observed for all tspans, and was more pronounced at the basal levels (Fig. 8C, brown *vs.* purple bars). These results suggested that in many cells, the release of EVs may be polarized on one side of the cell. One hypothesis is the existence of an activation of the transfer by specific acceptor cells. However, this hypothesis is not supported by the observation of spots deposited on the substratum in the absence of acceptor cells (Fig S3, lower panel, single basal plane). Another hypothesis is that the basal spots correspond to footprints, tspans-positive membranous structures left after migration^16,17^, an hypothesis supported by the pattern observed in some stacks of images (Fig. S3). To address this hypothesis cocultures of cells expressing mCherry-tagged CD9 or CD81 with CTB-labelled cells were imaged every 10 minutes at four different planes. As shown in Fig. 9, live events of transfer were observed, and showed mCherrry-CD81 spots released in upper planes from a donor cell that did not migrate during the release time frame (Fig 9A, open arrowheads), suggesting that migration is not the sole mechanism leading to transfer. Interestingly, the transfer was seen occurring all along the 12 hours timelapse, was oriented, and from a region of the plasma membrane below which we noticed a large fluorescent compartment (Fig. 9B, yellow arrowheads) which could be a supplying pool of cargoes for transfer. The presence of basal spots was observed in living cells co-cultures as shown in Fig. 9C for mCherry-CD9, with acceptor cells on top (Fig 9C, open arrowheads) or not (Fig. 9C, green arrowheads). Over time, the donor cell spreaded and covered the regions where basal spots were deposited (Fig 9C, yellow arrowhead). Altogether these data suggest that the cells released EVs in a polarized manner which is consistent with previous findings showing that truly polarized cells release EVs asymmetrically^18^.

**Figure 8:**
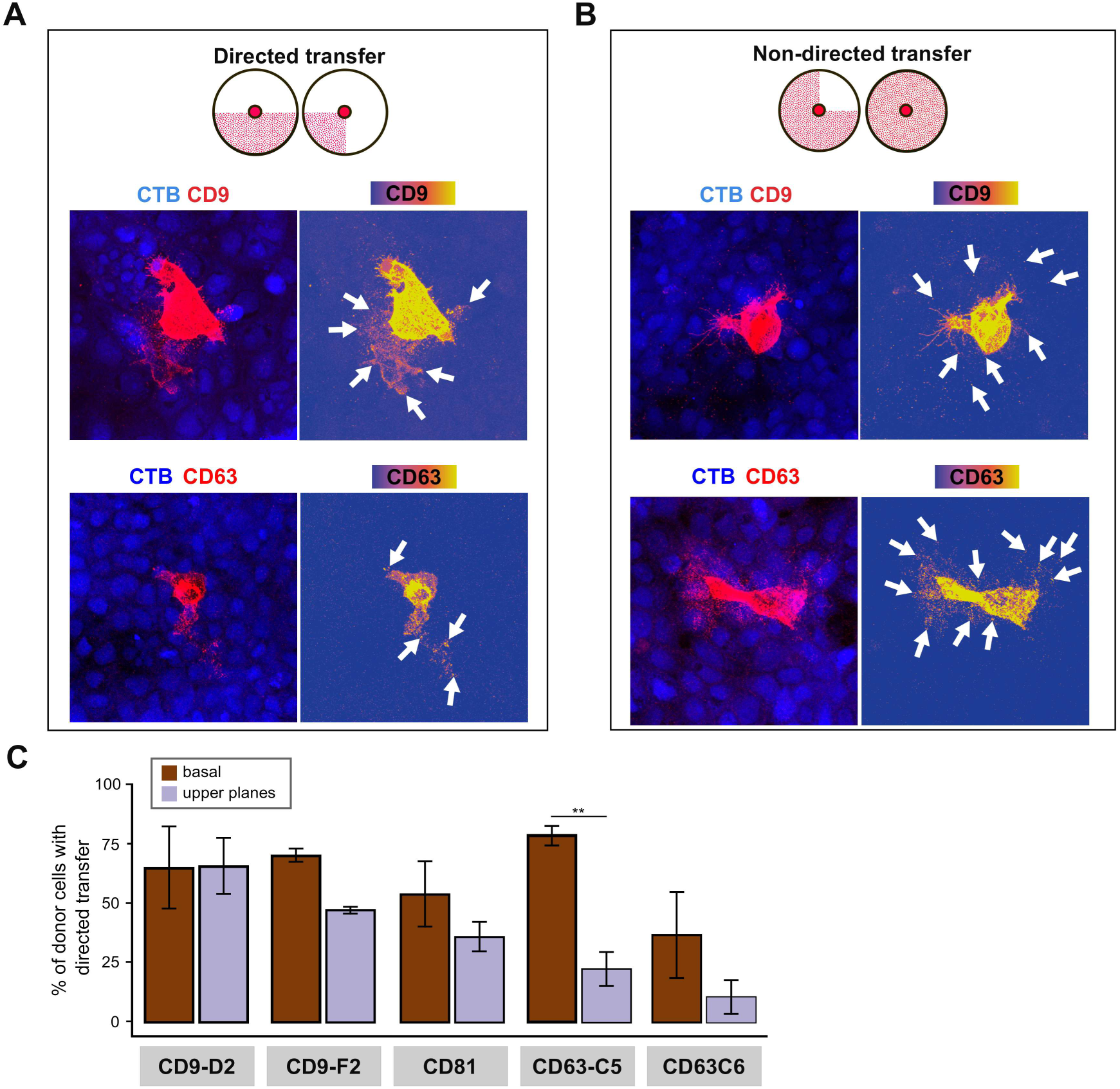
Transfer is directed. MCF-7 cells wt cells were cocultured for 48 h with MCF-7 cells KO for CD9, CD81, or CD63 stained with Cell Tracker Blue (CTB). Cells were fixed, permeabilized and immunostained against CD9, CD81, or CD63 revealed by a secondary antibody F(ab)^2^-AF555 prior imaging under a Leica SP5 confocal microscope. **A.** Representative maximal projections of stacks of images showing transfer directed to one side, or less. **B.** Representative maximal projections of stacks of images of transfer non-directed. **C.** Quantification of the percentage of donor cells with oriented transfer.

**Figure 9:**
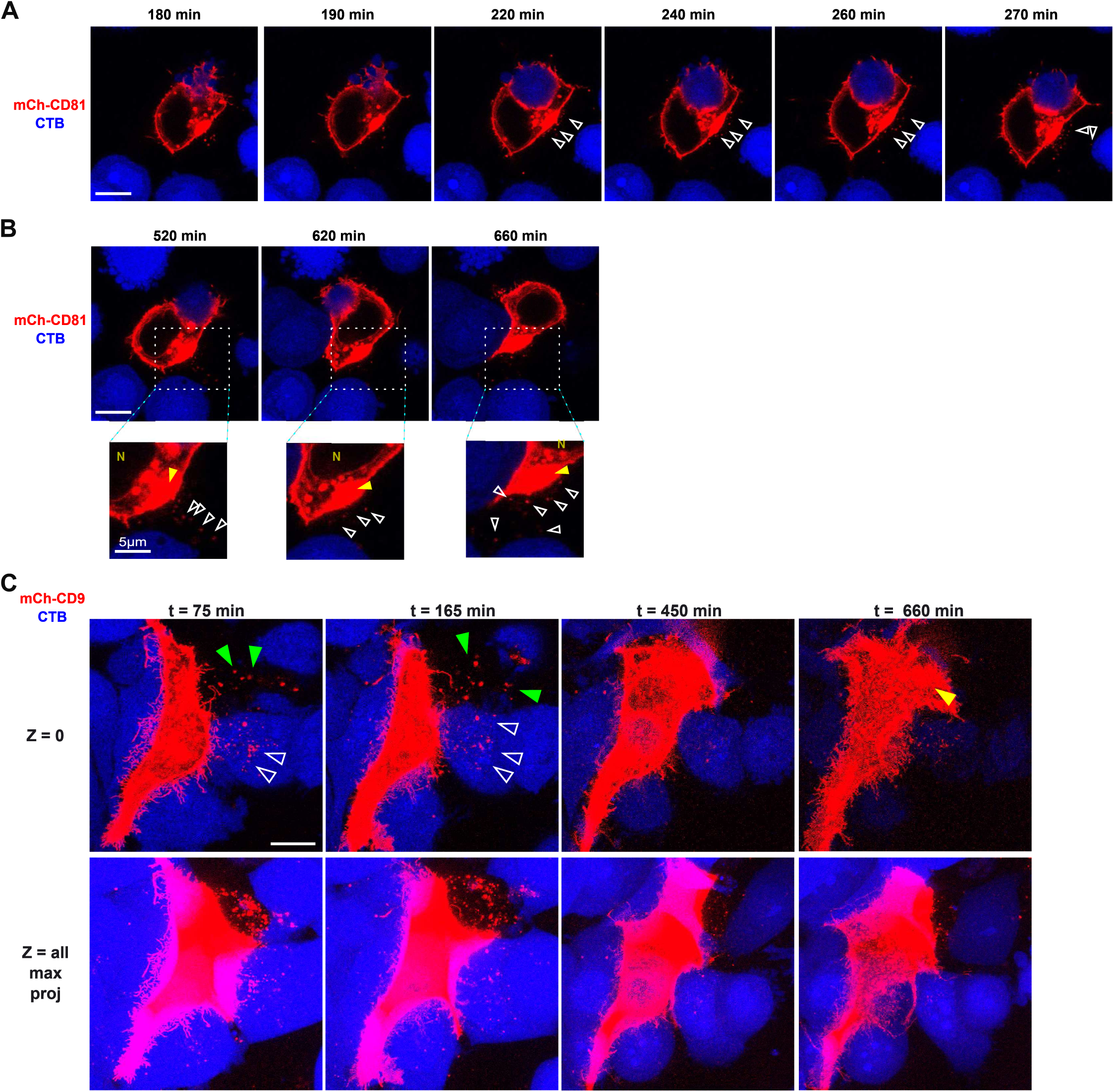
Live cell imaging of the transfer of mCherry-CD81 and mCherry-CD9. MCF-7 cells transiently expressing mCherry-CD9, mCherry-CD81 or mCherry alone were cocultured for 48 h with non-transfected MCF-7 cells stained with Cell Tracker Blue (CTG). **A-C.** Cells were imaged right after seeding, for 12 hours every ten minutes or every 15 minutes (C). Scale bars: 10 µm, unless indicated otherwise (1 µm in insets).

### Syntenin regulates the transfer of EV markers in coculture

As the biogenesis of CD63-containing EVs has been proposed to be regulated by the protein syntenin-1 in MCF-7 cells^10^, we investigated whether inhibition of syntenin-1 expression may affect the intercellular transfer of CD63, CD9 and CD81 in our system. To that end, wild type donor MCF-7 cells were transfected with siRNA control or against syntenin-1 as described elsewhere^10^, and cocultured with MCF-7 cells KO for CD9, CD81 or CD63. Depletion of syntenin in donor cells significantly decreased the transfer rate of all tspans (Fig. 10A) by a 4.4 and 2.7 fold change for CD63-C5 and C6 respectively, and a 1.9 and 2.5 for CD81 and CD9, respectively while the number of spots was decreased by 1.9 (CD63-C5/C6), 1.5 (CD9), and 3.3 (CD81)(Fig. 10B). Next, we investigated whether syntenin might regulate the 3-D topology of the transfer. To that end, we compared the distribution of spots along the (x,y) axis (Fig. 10C). The transfer still occurred mainly at low distance after syntenin depletion, with however a statistically significant decrease of the transfer to the clone CD63-C6 (Fig. 10C). Surprisingly depletion of syntenin caused a decrease in the number of spots at basal position for CD9 and CD63 (in the coculture with clone C6)(Fig. 10D-E). Thus, syntenin depletion may not only reduce EV production but may also reduce their ability to attach at basal position. In this regard, it has previously been demonstrated EVs purified from syntenin-depleted MCF-7 cells attached less efficiently to glass coverslips^6^ than control EVs.

**Figure 10:**
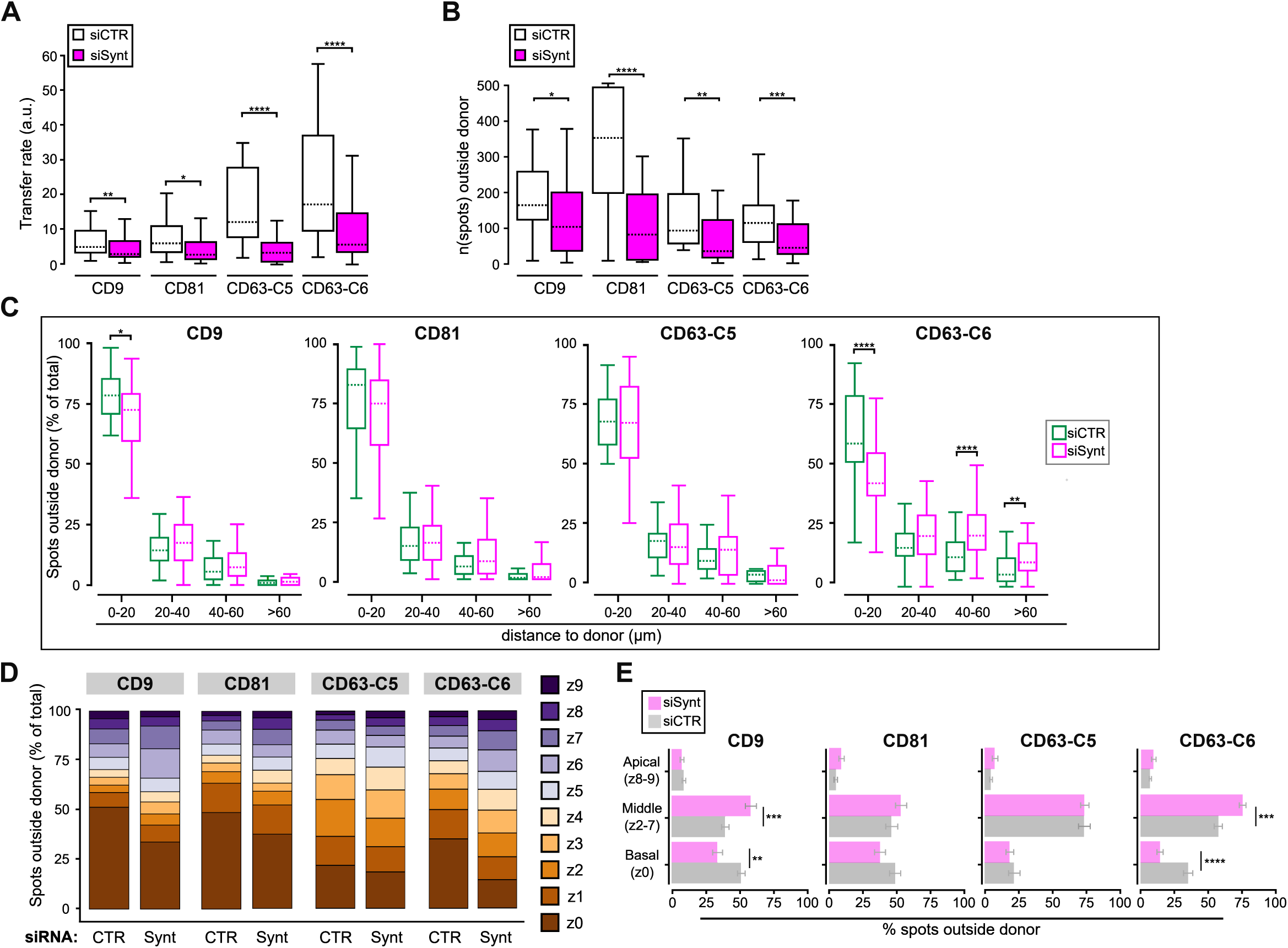
Effect of siRNA inhibition of expression of syntenin on the transfer of endogenous CD9, CD63 or CD81. Donor MCF-7 cells treated for 48 h with siRNA against syntenin (siSynt) or control (siCtrl) were cocultured for 48 h with acceptor MCF-7 cells KO for tspans CD9, CD81 and CD63 and stained with Cell Tracker Blue (CTB). Fixed and permeabilized cells were immunostained with primary antibodies against CD9, CD81 or CD63. **A.** Transfer rate. **B.** Number of spots detected outside donor cells. **C.** Percentage of spots at a given distance from donor cells (0-20 µm, 20-40 µm, 40-60 µm, >60µm) surrounding control cells (in green) or syntenin-1 depleted cells (in pink). **D-E.** Percentage of positive spots outside donor cells on individual confocal planes (z0–z9) (D), or on basal, summed middle and summed apical planes (E). Number of stacks analyzed: CD9: n= 37, CD81: n = 28, CD63-C5: n = 17, CD63-C6: n = 45 for siCTRL conditions, CD9: n = 32, CD81: n = 43, CD63-C5: n = 35, CD63-C6: n = 36 for siSynt conditions.

## Conclusions

Altogether, our results suggest that the transfer of EV markers is short distance, even in the presence of a flow. The distance transferred depended on the EVs marker considered, suggesting that different subclasses may travel more or less from donor cells, as those containing CD63 would travel further away than those containing CD9. Tracking single EVs in the three dimensions showed accumulation of excreted EV markers at the basal level, a localization that seemed to depend on the expression of syntenin in donor cells. Altogether, our work indicates that this system is suitable to study the topology and length of transfer of EV markers, endogenous and exogenous, and to find regulators of the intercellular transfer of EVs.

## Materials and Methods

### Cells, antibodies, plasmids and small interfering RNAs

The MCF-7 wild-type and knockout cells were generated and described previously^2^. Clones of CD63 and CD9 knockout MCF-7 cells were generated from mix of clones by limited dilution. Two clones were kept per cell line. The mCherry-CD9 and mCherry-CD81 constructs were generated by cloning CD9 and CD81 in a vector derived from pEGFP-C1 in which EGFP was replaced by mCherry. The primary antibodies used for immunofluorescence were mouse monoclonal antibodies against hCD9 (Ts9, IgG1), hCD81 (Ts81, IgG2a), and hCD63 (Ts63b, IgG2b) were described before^19,20^. CD81 antibody (clone 5A6) was purchased from ProteinTech, CD9 antibody (clone Mm2/57) from Biorad. For western blotting, we used a rabbit monoclonal antibody against syntenin-1 (Abcam, EPR8102) and a mouse monoclonal antibody against alpha adaptin subunit clone 100/2 (1:500) (SCB 58125). Secondary antibodies used were as follows: F(ab)2’-Goat anti-mouse Alexa Fluor Plus 555 (ThermoFisher A-48287) (1:1500), or Goat anti-Mouse IgG1 Alexa Fluor 568 (ThermoFisher A-21124) or Goat anti-Mouse IgG2b Alexa Fluor 488 (ThermoFisher A-21141) (1:500). The RNAi targeting human syntenin-1 sequence (siGenome SMART set of 4 oligo syntenin-1/SDCBP) and the non-targeting control (siStable non-targeting siRNA #1) were purchased from Dharmacon. A pool of 4 siRNAs sequences specific for human syntenin were used (GCAAGACCUUCCAGUAUAA, UAACAUCCAUAGUGAAAGA, GAAGGACUCUCAAAUUGCA, GGAUGGAGCUCUGAUAAAG) and the control sequence (UAGCGACUAAACACAUCAA) as described earlier^10^.

### Cell-culture and staining

MCF-7 cells were cultured in Dulbecco’s modified Eagle’s medium (DMEM) supplemented with 10% fetal bovine serum and 1% penicillin/streptomycin, at 37 °C in a humidified 5% CO₂ atmosphere. For coculture, cells were stained in culture medium with 20 µM CellTracker Green or CellTracker Blue (ThermoFisher, C2925, C2110) for 45 minutes to 1 hour at 37 °C. Cells were washed briefly with PBS (1X) and washed twice for 30 minutes in cell culture medium.

### Transfections

Cells at 40% confluency were transfected with INTERFERin (Polyplus, 101000028) with 50 nM siRNA (siControl or siSyntenin), according to the manufacturer’s instructions. Cells were trypsinized and used 48 h post siRNA transfection. Cells transfected by plasmids using FugeneHD (Promega, E2311) were trypsinized and used for coculture 24 h post plasmid transfection.

### Co-culture assay

Donor and acceptor cells were plated on poly-ornithine (Sigma-Aldrich, P4957) coated 1.5 H coverslips at 1:400 ratio. After 1 h in culture, medium was removed and replaced with medium supplemented with 2 mM 2’-deoxythymidine (dThymidine) (Sigma-Aldrich, T9250) as described earlier^9,11^. Cocultures of cells were done for 48 h at 37°C in a humidified, 5% CO_2_ atmosphere, unless stated otherwise. For cocultures under agitation, 24 h after dThymidine addition, plates were placed on a rocking shaker at 30 rpm speed inside the incubator for another 24 h. At the end of the coculture, cells were washed twice with ice-cold 1X PBS prior fixation for 30 min in ice cold 4% paraformaldehyde (PFA) on ice (EM grade, Delta Microscopies D15710). PFA was removed and cells were washed with 1X PBS before incubation with 50mM NH_4_Cl in PBS at room temperature for 15 min. For coculture with donor cells transfected with mCherry fusion proteins, Vectashield Vibrance media (Vector Laboratories, H-1700) was used to mount cells. For endogenous transfer assay, coculture of wild-type and KO cells were washed with 1X PBS prior incubation for 1 h with primary antibodies in staining buffer (1% bovine serum albumin (BSA), 0.1% saponin in 1X PBS). Coverslips were washed thrice 5 min in staining buffer before incubation with secondary antibodies in staining buffer for 1 h at room temperature. Coverslips were then rinsed in 1X PBS 1% BSA, mounted using Vectashield Vibrance mounting media supplemented or not with DAPI (Vector Laboratories, H-1800).

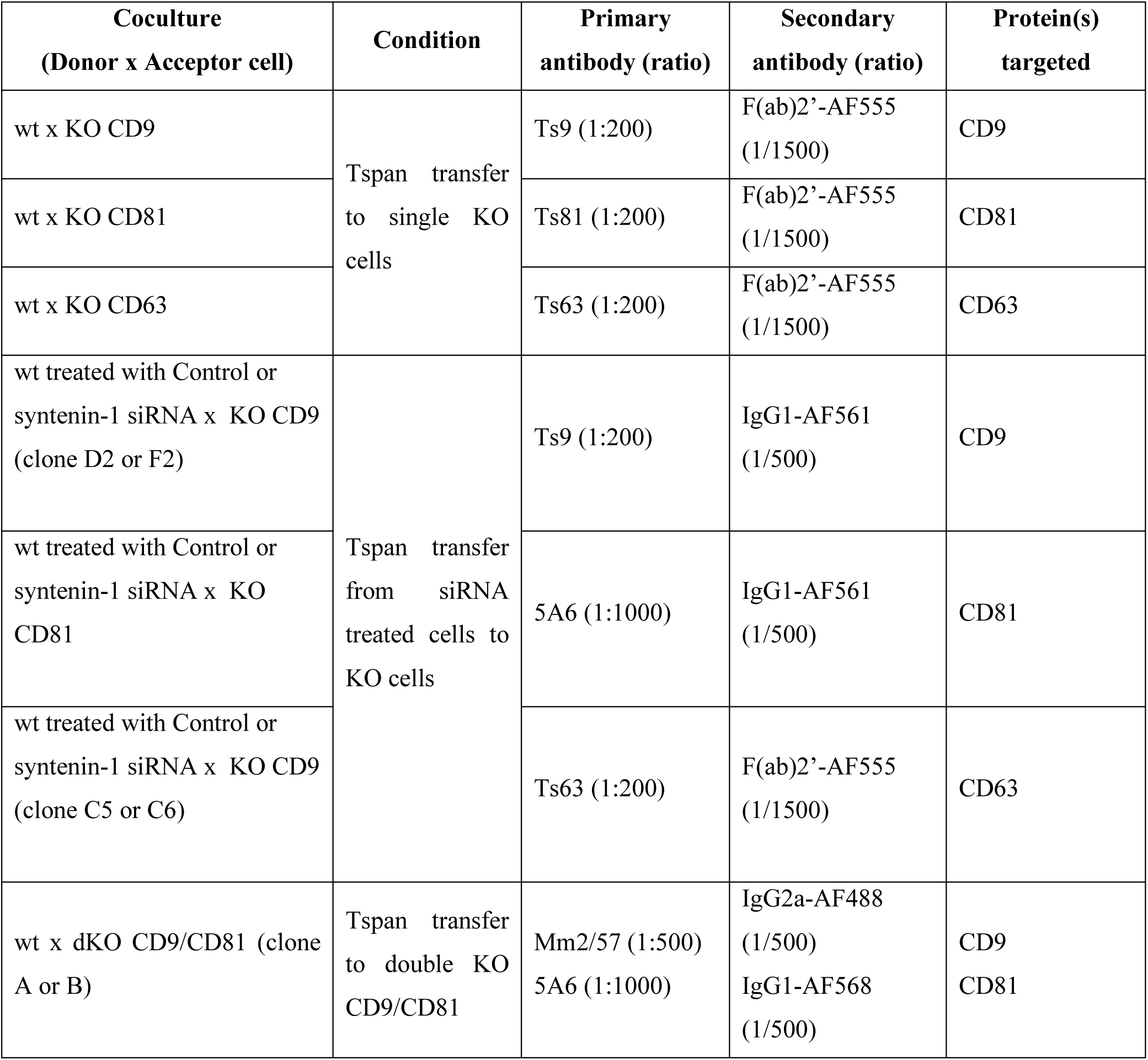

## Imaging

Confocal microscopy was performed on a Leica TCS SP8 at the INRAE MiMA2 facility (Jouy-en-Josas, FR) with a 63X, 1.2 NA, objective. For quantification, stacks of images were acquired with the following parameters: 1.28 zoom, 700Hz, 1024×1024, 16 bits. When CTB was used, imaging was performed on a Leica TCS SP5 at Necker SFR imaging facility (Paris, FR) with a 40X 1.25 NA objective, 2.5 zoom, 1000Hz, 512×512, 16 bits. Stacks were acquired provided the following criteria : presence of donor cell, confluency, absence of dead cells or large debris, distance from at least 8 cell diameters to the closest donor cells. Distant regions were regions of the coculture containing only acceptor cells, distant from at least 8 cells diameter to the closest donor cell. The laser power and PMT voltage were adjusted for each experiment but kept identical within experiment, and set to detect weak signals in acceptor cells without saturation of donor cells, as described earlier^9,11^. Cells were sampled in 10 planes covering the entire cellular height, from basal to apical.

### Automated image analysis for transfer quantification

Donor cells were detected automatically on each plane by the cell shape plugin { https://doi.org/10.5281/zenodo.4317782} of ICY software { <u>10.1038/nmeth.2075</u>} and parameters of donor cells acquired (volume, fluorescence intensities). The 3D ROI corresponding to the donor cell was cut off after being enlarged by 4 pixels in the (x,y) direction to remove donor cell membrane that may generate falsely be accounted for transfer. Quality control on each stack was done to ensure correct contouring of donor cell and manual correction was applied if protrusions were not removed by the enlargement. To quantify the transfer in distant regions in which no donor cells were present but only acceptor cells were present, we used a representative donor ROI from the corresponding experiment to mimick the presence of a donor cell. To normalize the transfer in these regions, *i.e.* surrounding this “artificial” ROI, we used the average fluorescence intensity value of donor cells in the corresponding experiment. For each stack, the transfer intensity outside donors was calculated as the sum fluorescence intensity outside donor normalized to that inside donor. Automated spot detection was done on ICY software. Stacks of images were subjected to Gaussian filtering. The “distant” regions devoid of donor cells were used to find the best threshold for transfer quantification on fields with donor cells. For endogenous cocultures, the maximum entropy thresholding method was applied in distant regions, therefore specific to each experiment. For transient coculture experiments, the K-means thresholding method (classes 2 to 4, depending on the images) was used. In these cases, distant regions were thresholded based on the average K-mean threshold value obtained from the donor object regions. Fluorescent pixels above the threshold value and located outside the donor were considered as positive spots using a wavelet spot detector. Spots z-axis position and sum fluorescence intensity were measured in each channel. The shortest distance to the donor object contour was calculated using the Distance Transform Chamfer 5 algorithm on ICY software, altogether generating the (x,y,z) coordinates for each spot, along with its fluorescence intensity. For each stack, the transfer rate was calculated as the sum fluorescence in spots normalized to the sum fluorescence in donor cell.

### Statistical analysis

Normality was assessed using the Shapiro–Wilk test. For non-normal data, the Wilcoxon rank-sum (independent samples) or Wilcoxon signed-rank (paired samples) test was used. Normally distributed data were analyzed with two-tailed Student’s t-tests. Average +/- sem if not indicated otherwise. All statistical analysis were performed on RStudio (version 2024.09.0+375){Posit team (2024). RStudio: Integrated Development Environment for R. Posit Software, PBC, Boston, MA. URL http://www.posit.co/.} and p < 0.05 (*), p < 0.01 (**), p < 0.005 (***), p < 0.001 (****) were considered statistically significant.

## Acknowledgments and funding

We acknowledge the French National Research Agency (ANR) research grant ANR-21-CE34-0007 NanoMilk (A.B.)

---

**Supplementary Fig. S1**

**A.** Percentage of donor cells with transfer. **B**. Average mCherry expression level in donor cells. **C-D.** Number of mCherry-positive spots for each donor cell plotted against donor cell intensity (C) or donor cell volume (D).

**Supplementary Fig. S2**

**A.** Number of CD9-, or CD81-, or CD63-positive spots for each donor cell plotted against donor cell volume or intensity. **B**. Percentage of donor cells with transfer. **C**. Percentage of spots containing CTB. **D.** Average number of spots per plane.

**Supplementary Fig. S3**

**A.** Images of MCF-7 cells wt cocultured with KO cells for CD81 or CD63 that were CTB stained prior coculture. Arrows point to footprints-like structures. Scale bars: 20µm.

## References

1. van Niel, G., D’Angelo, G. & Raposo, G. Shedding light on the cell biology of extracellular vesicles. Nat. Rev. Mol. Cell Biol. 19, 213–228 (2018).

2. Fan, Y., et al. Differential proteomics argues against a general role for CD9, CD81 or CD63 in the sorting of proteins into extracellular vesicles. J. Extracell. Vesicles 12, e12352 (2023).

3. Jeppesen, D. K. et al. Reassessment of Exosome Composition. Cell 177, 428–445.e18 (2019).

4. Mathieu, M. et al. Specificities of exosome versus small ectosome secretion revealed by live intracellular tracking of CD63 and CD9. Nat. Commun. 12, 4389 (2021).

5. Nigri, J. et al. CD9 mediates the uptake of extracellular vesicles from cancer-associated fibroblasts that promote pancreatic cancer cell aggressiveness. Sci. Signal. 15, eabg8191 (2022).

6. Irmer, B., et al. Syntenin Controls Extracellular Vesicle-Induced Tumour Migration by Regulating the Expression of Adhesion Proteins on Small Extracellular Vesicles. J. Extracell. Vesicles 14, e70133 (2025).

7. Tognoli, M. L. et al. Lack of involvement of CD63 and CD9 tetraspanins in the extracellular vesicle content delivery process. *Commun*. Biol. 6, 532 (2023).

8. Verweij, F. J. et al. Live Tracking of Inter-organ Communication by Endogenous Exosomes In Vivo. Dev. Cell 48, 573–589.e4 (2019).

9. Burtey, A. et al. Intercellular transfer of transferrin receptor by a contact-, Rab8-dependent mechanism involving tunneling nanotubes. FASEB J. 29, 4695–4712 (2015).

10. Baietti, M. F. et al. Syndecan–syntenin–ALIX regulates the biogenesis of exosomes. Nat. Cell Biol. 14, 677–685 (2012).

11. Frei, D. M. et al. Novel microscopy-based screening method reveals regulators of contact-dependent intercellular transfer. Sci. Rep. 5, 12879 (2015).

12. de Chaumont, F. et al. Icy: an open bioimage informatics platform for extended reproducible research. Nat. Methods 9, 690–696 (2012).

13. Fordjour, F. K., Guo, C., Ai, Y., Daaboul, G. G. & Gould, S. J. A shared, stochastic pathway mediates exosome protein budding along plasma and endosome membranes. J. Biol. Chem. 298, 102394 (2022).

14. Grangier, A., Wilhelm, C., Gazeau, F. & Silva, A. High yield and scalable EV production from suspension cells triggered by turbulence in a bioreactor. Cytotherapy 22, S50 (2020).

15. Gazeau, F., Silva, A. K. A., Merten, O.-W., Wilhelm, C., Piffoux, M. Fluid system for producing extracellular vesicles and associated method. (2019).

16. Yamada, M. et al. Substrate-attached materials are enriched with tetraspanins and are analogous to the structures associated with rear-end retraction in migrating cells. Cell Adhes. Migr. 7, 304–314 (2013).

17. Perez Ipiña, E., d’Alessandro, J., Ladoux, B. & Camley, B. A. Deposited footprints let cells switch between confined, oscillatory, and exploratory migration. Proc. Natl. Acad. Sci. 121, e2318248121 (2024).

18. Colombo, F. et al. Polarized cells display asymmetric release of extracellular vesicles. Traffic 22, 98–110 (2021).

19. Charrin, S. et al. Rapid Isolation of Rare Isotype-Switched Hybridoma Variants: Application to the Generation of IgG2a and IgG2b MAb to CD63, a Late Endosome and Exosome Marker. Antibodies 9, 29 (2020).

20. Charrin, S. et al. The Major CD9 and CD81 Molecular Partner: IDENTIFICATION AND CHARACTERIZATION OF THE COMPLEXES*. J. Biol. Chem. 276, 14329–14337 (2001).

