## Supplementary figures and images for "The intercellular transfer of extracellular vesicles markers CD63, CD9 and CD81 is spatially polarized and restricted to cell vicinity"

### SimonMG Supp.pdf

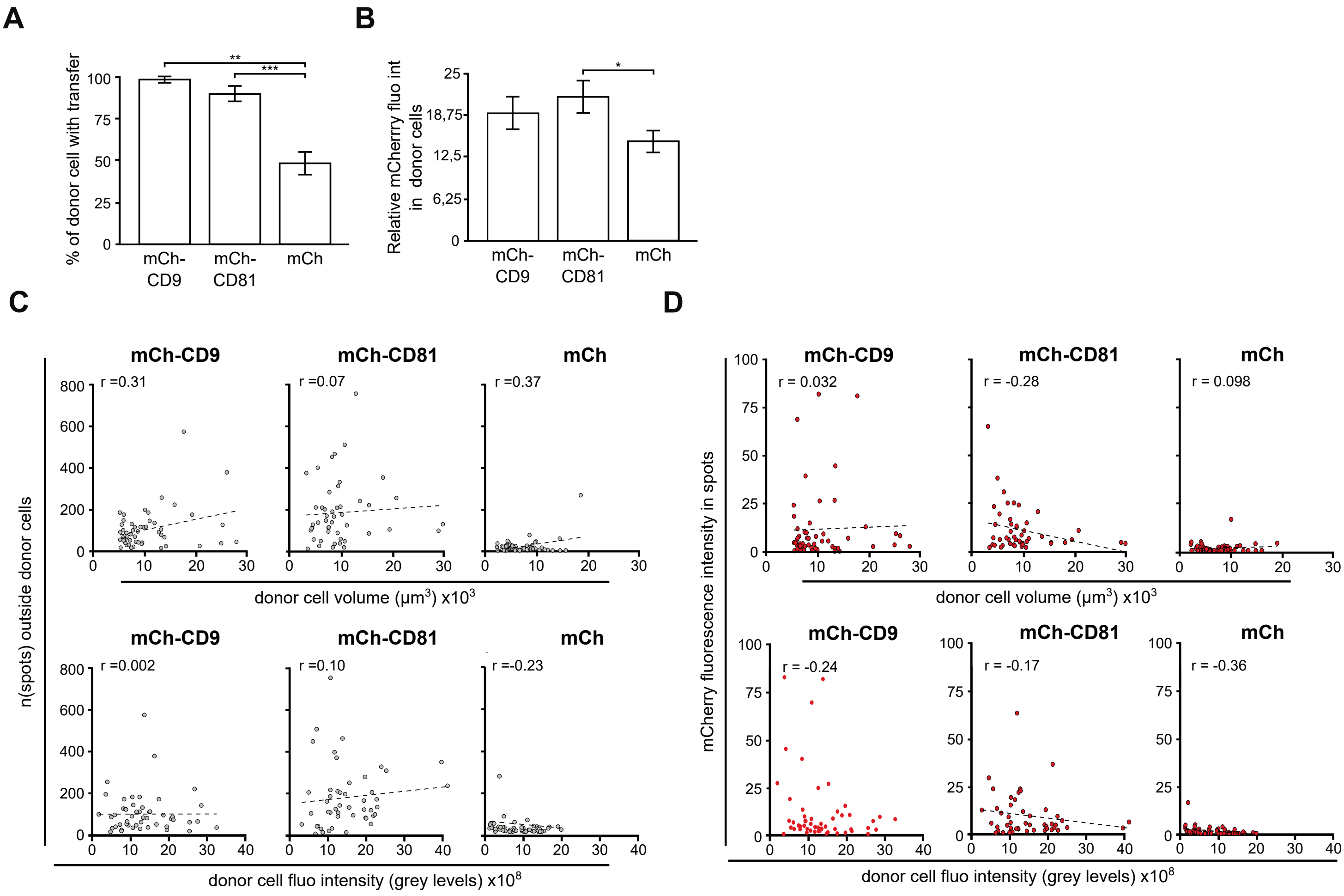

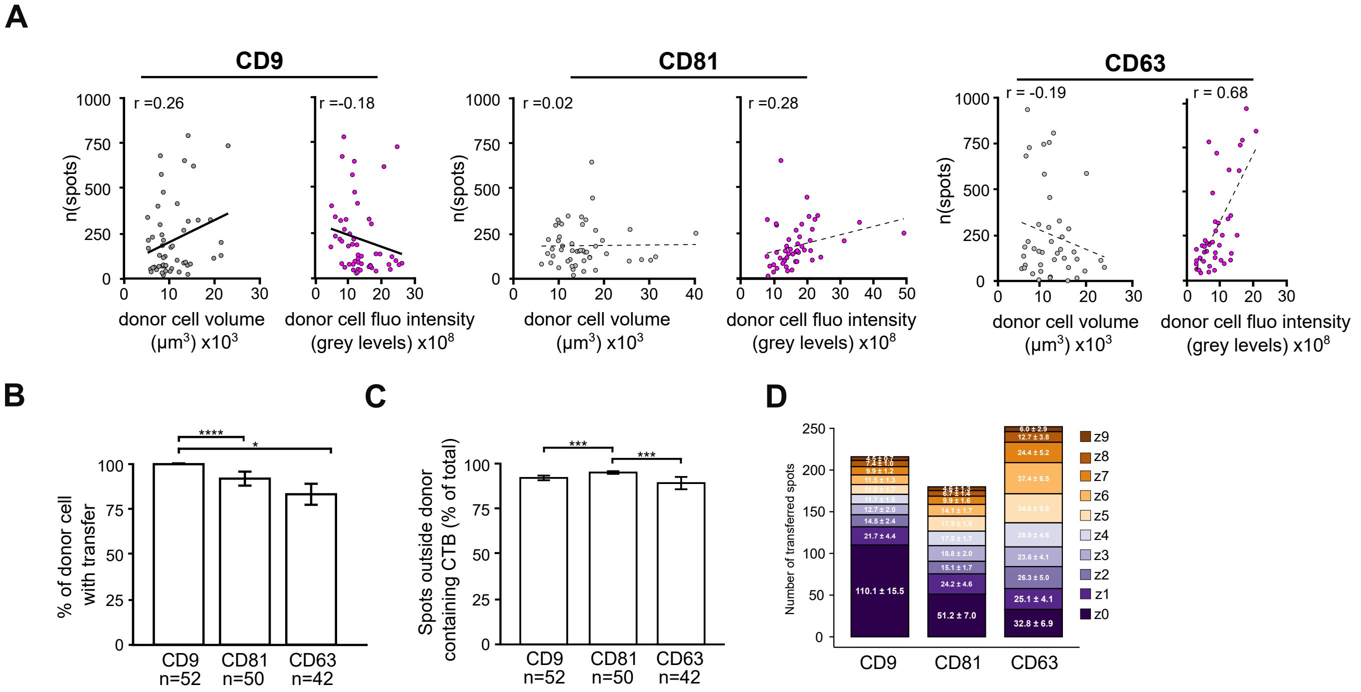

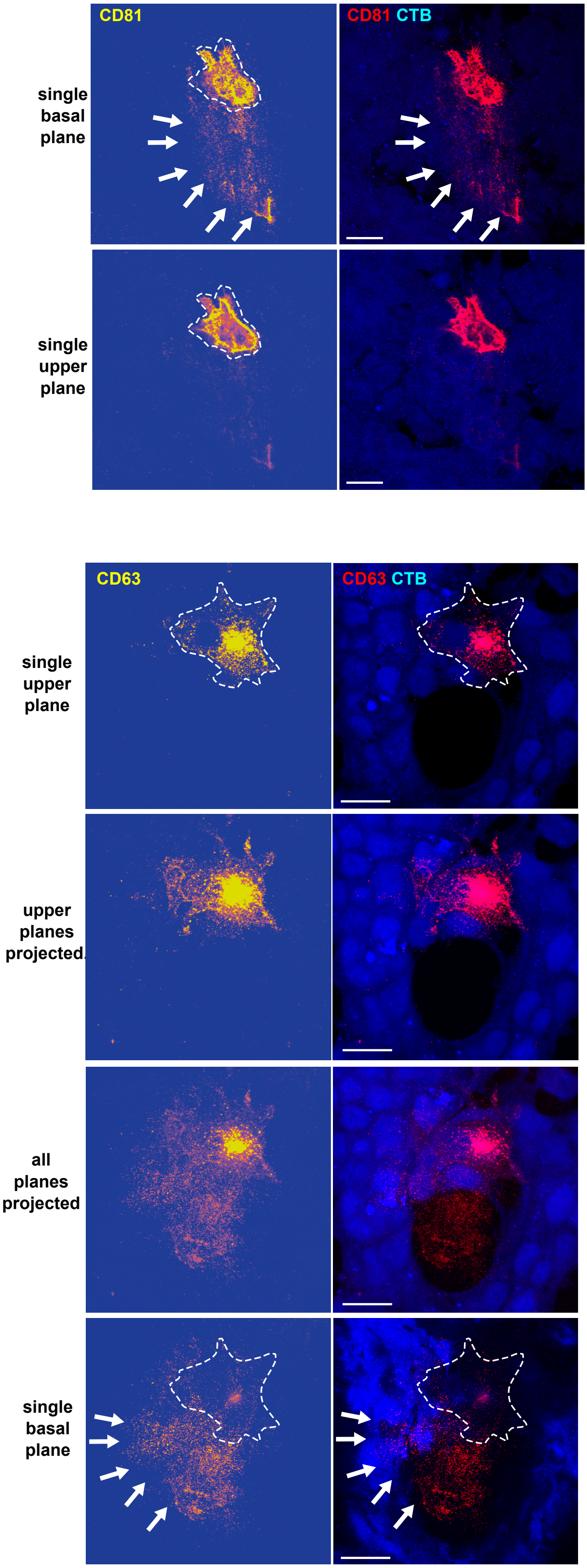
